# Approaches by *Rhodotorula mucilaginosa* from a chronic kidney disease patient for elucidating the pathogenicity profile by this emergent species

**DOI:** 10.1101/701052

**Authors:** Isabele Carrilho Jarros, Flávia Franco Veiga, Jakeline Luiz Corrêa, Isabella Letícia Esteves Barros, Marina Cristina Gadelha, Morgana F. Voidaleski, Neli Pieralisi, Raissa Bocchi Pedroso, Vânia A. Vicente, Melyssa Negri, Terezinha Inez Estivalet Svidzinski

**Author notes:** **Corresponding author**: Terezinha Inez Estivalet Svidzinski, Division of Medical Mycology, Teaching and Research Laboratory in Clinical Analysis – Department of Clinical Analysis of State University of Maringá – Paraná – Brazil, Av. Colombo, 5790, CEP: 87020-900, Maringá, PR., Brazil, or.

## Abstract

**Background:** Traditionally known as a common contaminant, *Rhodotorula mucilaginosa* is among the leading causes of invasive fungal infections by non-candida yeasts. They affect mainly immunocompromised individuals, often mimicking the cryptococcosis infection, despite invasive infections by *Rhodotorula* are still not well explained. Thus, here we aimed to characterize microbiologically clinical isolates of *R. mucilaginosa* isolated from colonization of a patient with chronic renal disease (CKD), as well as to evaluate their phylogeny, antifungal susceptibility, virulence, and pathogenicity in order to infer the potential to become a possible infection.

**Methodology/Principal Findings:** For this study, two isolates of *R. mucilaginos*a from oral colonization of a CKD patient were isolated, identified and characterized by classical (genotypic and phenotypic) methods. Susceptibility to conventional antifungals was evaluated, followed by biofilm production, measured by different techniques (total biomass, metabolic activity, colony forming units and extracellular matrix quantification). Finally, the pathogenicity of yeast was evaluated by infection of *Tenebrio molitor* larvae.

All isolates were resistant to azole and sensitive to polyenes and they were able to adhere and form biofilm on the abiotic surface of polystyrene. In general, similar profiles among isolates were observed over the observed periods (2, 24, 48 and 72 hours). Regarding extracellular matrix components of biofilms at different maturation ages, *R. mucilaginosa* was able to produce eDNA, eRNA, proteins, and polysaccharides that varied according to time and the strain. The death curve *in vivo* model showed a large reduction in the survival percentage of the larvae was observed in the first 24 hours, with only 40% survival at the end of the evaluation.

**Conclusions/Significance:** We infer that colonization of chronic renal patients by *R. mucilaginosa* offers a high risk of serious infection. And also emphasize that the correct identification of yeast is the main means for an efficient treatment.

**Author Summary:** The genus *Rhodotorula* is known to be a common contaminant, however, it has been increasing in the last years, reports of different forms infections by this yeast, reaching mainly individuals with secondary diseases or with low immunity. However, very little is known about the mechanism that triggers the disease. Thus, this study aims to characterize microbiologically clinical isolates of *R. mucilaginosa* isolated from a patient with chronic renal disease, as well as to evaluate their phylogeny, antifungal susceptibility, virulence, and pathogenicity in order to infer the potential to become a possible infection. It was possible to characterize in general the clinical isolates, to determine that they are resistant to an important class of the antifungal agents which are the azoles. In addition, they are able to adhere and to form biofilm on abiotic surfaces, this skill represents an important factor of virulence, which would guarantee their presence in medical devices, such as catheters, surfaces. These biofilm works as true reservoirs of these fungi disseminate and cause serious infections. This pathogenic potential was reinforced by a great reduction of survival in the larvae infected with this yeast. Therefore our results infer a high risk of infection to patients who are colonized by *R. mucilaginosa*.

## Introduction

Human fungal infections by *Rhodotorula* spp. are increasingly in the last decades [1], in China it is among the main cause of invasive fungal infections by non-candida yeasts [2] and is considered an emerging pathogen. It was classified as the third most commonly isolated yeast from blood cultures and the most common microorganism isolated from the hands of hospital employees and patients [3].

Species from this genus have been considered an opportunistic pathogen since they affect mainly immunocompromised individuals. A recent systematic review [1] shows fungal infections by *Rhodotorula* spp. consists mainly of bloodstream infections, as well as central nervous system (CNS), affecting especially patients under the use of central venous catheters (CVC). However, there are reports at the literature proving that *Rhodotorula* spp. causes besides fungemia and meningitis, also cutaneous infections, peritonitis, keratitis, ventriculitis, ocular and other less frequent infections [4–6]. These clinical presentations, as well as microbiological aspects, makes *Rhodotorula* infections look like cryptococcosis and to worsen this scenario, yeasts of this genus are usually azoles resistant, including isavuconazole, which has good *in vitro* activity against *Cryptococcus* species, but it is not effective against *Rhodotorula* spp. [7]. Reinforcing that the management of these two fungal infections needs to be well known.

Although, endocarditis by *Rhodotorula* spp. has already been related to immunocompetent patients [8, 9] apparently without risk factor for opportunistic infection, invasive infections caused by this yeast are mainly associated with underlying immunosuppression. The most affected are patients with severe diseases such as leukemia, cancer or other solid tumors, lymphoproliferative disease, HIV, diabetes mellitus and submitted to various types of surgery [10–13]. Recently has been described some cases of endocarditis by *R. mucilaginosa* in chronic kidney disease (CKD) patients [9,14,15]. Nevertheless, despite their importance, invasive *Rhodotorula* infections are not well explained yet, few is known about the virulence potential of this yeast, and whether the dissemination process depends more of pathogenic merit of the fungus or the patient debility. In fact, CKD has become a serious public health problem, affecting 8 to 16% of the world’s population, it is responsible for high morbidity and mortality and represents a heavy financial burden for public health systems in developing and developed countries [16]. CKD implies in structural and functional renal deficiencies, which result in complex disorders. There is a close relationship between the progressively defective immune system with side effects, including cardiovascular problems, infections and malignancies [17].

The association of *Rhodotorula* with CVC or other invasive medical devices is justified by its ability of biofilm production [1]. These devices provide appropriate surfaces for biofilms formation and establishment. However, in contrast to the extensive literature on biofilms of *Candida* species, little attention has been paid to emerging fungal pathogens, such as *Rhodotorula* species [18, 19].

The biofilm installation at some surfaces contacted to the host can trigger an acute fungemia and consequently a disseminated infection. This occurs when the clusters of cells are dispersed from the initial biofilm and occupy a not colonized niche [20]. Recent studies have shown that cells that detach themselves from a biofilm have a greater association with severe infections, with high mortality rates compared to microorganisms in their planktonic form. In fact, more than 65% of human infections involve the formation of biofilms, keeping up to the growing number of immunocompromised patients. In addition, more than 500,000 deaths per year are caused by biofilm-associated infections [21]. However, little is still known on *R. mucilaginosa* from mucosa colonization of immunocompromised CKD patients.

*Rhodotorula* yeasts are part of the human microbiota as commensal microorganisms of skin, nails, gastrointestinal, urinary, and respiratory tracts. They are also widely found in nature and has been isolated from environmental sources, like air, soil, and plants [22]. Falces-Romero *et al*. 2018 isolated *R. mucilaginosa* from blood cultures of eight patients, six of them had a real infection and two were considered contaminants, since usually the yeasts causing infection, are from environmental origin or even from the microbiota itself [4–6]. As an opportunistic agent, it is indeed not easily differentiated from colonization or infection. Recently, we proved that *R. mucilaginosa* was able to colonize and crossed a device used for dermis regeneration of burned patients in three days increasing significantly in seven days, therefore offering high risk for systemic infection [23]. In that occasion, we demonstrated that commensal yeasts, commonly found at the environment, skin or mucosa of health professionals and patients, could offer a risk of infection for severe patients. In order to improve the knowledge about this question, we intend to expand the study addressed just to *R. mucilaginosa.* Thus, here we aimed to characterize microbiologically clinical isolates of *R. mucilaginosa* isolated from colonization of chronic renal patients as well as to evaluate some of the aspects related to phylogeny, antifungal susceptibility, virulence, and pathogenicity in order to infer the potential to become a possible infection.

## Materials and Methods

### 1.0 Studied group and isolation

For the purpose of this study, a patient was aleatorily selected from a bigger project involving 243 patients with CKD under the care of the nephrology service of a reference hospital in the northwest of Parana State, Brazil, between October and November 2014. This voluntary is a man, 55 years old, diabetic (*Diabetes Melittus* type II), confirmed with chronic kidney diseases (stage 5) at 2 years before, and he was under hemodialysis for 6 months, no using antifungals and absence of oral lesions. The data collection, the oral mucosa examination, were performed according to Pieralisi et al., 2016. This study was conducted according to the Resolution 466/2012 of the National Health Council and was previously approved by the Ethics Committee for the Research Involving Humans of the State University of Maringá, Brazil [COPEP-EMU n° 383979, CAEE resolution n° 17297713.2.0000.0104].

### 2.0 Microorganisms

This study was conducted with two clinical isolates from colonizations oral of CKD patients plus the *R. mucilaginosa* ATCC 64684. For the clinical isolates the collecting biological samples and the cultivation method were performed as described previously [24]. Briefly, yeasts were sub cultured in chromogenic medium CHROMagar™ *Candida* (Difco, USA), to check the culture purity. After, the isolates were identified by classical tests, including macro and micro morphologies, fermentation tests and assimilation of carbohydrate and nitrogen sources [25, 26]. To confirm the identification, mass spectrometry assisted by flight time desorption/ionization matrix (MALDI TOF-MS) was performed. For the MALDI TOF-MS method, the yeasts were prepared according to specific protocols [27, 28] with a Vitek MS mass spectrometer using the Myla or Saramis software for data interpretation.

These yeasts were deposited at Microbial Collections of Paraná Network-TAX online and on Micoteca of the Medical Mycology laboratory, Laboratório de Ensino e Pesquisa em Análises Clínicas of Universidade Estadual de Maringá (LEPAC), with the identification codes: CMRP3462 (MK453051; Genbank) and CMRP3463 (MK453052; Genbank). On LEPAC, the yeasts were stored in Sabouraud Dextrose Broth (SDB; Difco™, USA) with glycerol at –80 °C. All samples were cultured on SDA with additional chloramphenicol (0.1%) and incubated at 25 °C for up to 3 days, after all tests [29].

### 3.0 Morphological characterization

The morphology was assessed with by optical microscopy (EVOS™ FL, Life Technologies) and by Scanning Electron Microscopy (SEM; Quanta 250™, ThermoFisher). The colony, cell morphology and the polysaccharide capsule were observed by light microscopy at 40x magnification. The colony was observed after microcultive and analyzed directly by light microscopy [26]. To analyze capsule, a suspension of 500 µg/mL phosphate-buffered saline 0.01 mol/L, pH 7.4 solution (PBS) with one isolate colony and 500 µL of China ink was prepared and placed on a slide for observation under light microscope. The cell morphology was performed with Calcofluor White (Fluka Analytical, Canada) diluted in a proportion of 1:4 in PBS for 5 minutes, and excess dye was removed by washing once with PBS. The yeasts were observed with a filter capable of detecting the yeast cell wall (BP 365– 370, FT 400, LP 421). SEM analysis was performed at Laboratory of Electron Microscopy and Microanalysis, Universidade Estadual de Londrina, Londrina, Paraná, Brazil, supervised by Admilton G. de Oliveira, according to Negri et al., 2011, The samples were observed at 5000×magnification.

### 4.0 Genotypic characterization (sequencing and phylogenetic study)

The DNA extraction of isolates was performed according Vicente et al., (2008) [31], using a silica: celite mixture (silica gel H, Merck 7736, Darmstadt, Germany/Kieselguhr Celite 545, Machery, Düren, Germany, 2:1, w/w). The internal transcribed region (ITS) was amplified using the universal primers ITS1 (5’-TCCGTAGGTGAACCTGCGG-3’) and ITS4 (5’-TCCTCCGCTTATTGATATGC-3’) [31, 32]. PCR was performed in a 12,5 μL volume of a reaction mixture containing 4,3 μL of mix solution containing 0,3 mM dNTPs, 2,5 mM MgCl2, 1,25 μL reaction buffer, 0,5 μL of each primer (10 pmol) and 1 μL rDNA (20 ng/μL). The sequencing was performed by sanger method in automated sequencer ABI3730 (Applied Biosystems Foster City, U.S.A). Consensus sequences of the ITS region were inspected using MEGA v.7 software and alignments were performed using MAFFT interface (online). The identification of specie was determined by phylogenetic analysis, using type strains established by Wang et al., 2015 and Nunes et al., 2013. Phylogenetic tree was performed MEGA v.7 software with 1,000 bootstrap replicates using the maximum likelihood function and the best evolutionary model corresponding to the data set used. Bootstrap values equal to or greater than 80% were considered statistically significant.

### 5.0 Antifungal susceptibility profile

The antifungal susceptibility profile of *R. mucilaginosa* isolates was determined against amphotericin B (Sigma, USA), fluconazole (Sigma, USA), voriconazole (Sigma, USA), itraconazole (Sigma, USA) and nystatin (Sigma, USA). The test was performed by a microdilution assay in broth, according to the Clinical Laboratory Standards Institute (CLSI, 2008), M27-A3 document (CLSI, M27-A3). Concentrations ranged between 0.125 and 64 µg/mL for fluconazole, between 0.03 and 16 µg/mL for amphotericin B and voriconazole, between 0.0625 and 32 µg/mL for itraconazole and 0.25 and 128 µg/mL for nystatin. Suspensions were tested with antifungal solutions in 96-well microplates (Nunclon Delta; Nunc) incubated for 48 hours at 25 °C. *C. albicans* ATCC® 90028 was used as a control and the reading of microplates was performed at 405 nm (Expert Plus Microplate Reader; ASYS). The Minimum Inhibitory Concentration (MIC) was determined according to CLSI, M27-A. Results were given as: susceptible (S); susceptible dose dependent (SDD); and resistant (R). Cut-off points were: S ≤8 *μ*g/mL; SDD = 16–32 *μ*g/mL; and R ≥64 *μ*g/mL for fluconazole, S ≤ 1 *μ*g/mL; SDD = 2 *μ*g/mL; R ≥16 *μ*g/mL for voriconazole, S ≤0.125 *μ*g/mL; SDD 0.25-0.5 *μ*g/mL; R ≥32 *μ*g/mL for itraconazole, S≤ 4 *μ*g/mL; SDD = 8–32 *μ*g/mL; R ≥ 64 *μ*g/mL for nystatin. For amphotericin B, resistant isolates were defined as isolates with MIC >1 µg/mL [34].

### 6.0 Adhesion and Biofilm formation assays

The biofilm formation assay was adapted from previously described method [19]. The strains initially cultured in SDA at 25 °C for 72 hours were further subcultured into SDB and grown for 18 hours with shaking at 110 rpm at 25 °C. The grown cultures were harvested, washed twice with phosphate-buffered saline (PBS; pH 7.2), and adjusted to a concentration of 1×10^7^ cells/mL, using a Neubauer chamber in RPMI 1640 medium (Roswell Park Memorial Institute, Gibco). Biofilm formation was tested in sterile 96-well polystyrene flat-bottom plates (TPP®, Trasadingen, Switzerland) with 200 μL of inoculum. A test medium without yeasts was performed and used as a negative control. The plates were then incubated with agitation at 110 rpm at 25 °C for 2 hours. After this time, the supernatant was gently removed from the wells, the cells were washed three times with PBS for removal of non-adherent yeasts, 200 μL of RPMI 1640 were added and the plates were incubated at 110 rpm at 25 °C for 24, 48 and 72 hours. The processes of biofilm formation was evaluated after the adhesion (2 hours) and the different ages of biofilm maturation (24, 48 and 72 hours).

#### 6.1 Biofilm characterization

The adhesion and biofilms were analyzed by number of cultivable cells determined by counting colony forming units (CFU); metabolic activity by the tetrazolium salt 2,3-bis(2- methoxy-4-nitro-5-sulfophenyl)-5-(phenylamino)-carbonyl-2H-tetrazoliumhydroxidand (XTT; Sigma-Aldrich, USA) reduction assay; total biofilm biomass by crystal violet staining (CV); quantification of proteins, polysaccharides, extracellular DNA (eDNA) extracellular RNA (eRNA) in biofilm matrix using spectrophotometer; and finally biofilm structure by SEM.

Briefly, the wells with different ages of biofilm (2, 24, 48 and 72 hours) were washed twice in PBS to remove loosely attached cells. Before determining the CFU, the time and potency of sonication were optimized in order to allow the complete removal of the adhered cells without causing any damage. After washing each well, the biofilms were resuspended with 100 μL of PBS and scraped. The suspensions were transferred a new tub and sonicated (Sonic Dismembrator Ultrasonic Processor, Fisher Scientific) for 50 seconds at 30%, and then the suspension was vortexed vigorously to disrupt the biofilm matrix and serial decimal dilutions, in PBS, were plated onto SDA. Agar plates were incubated for 48 hours at 25 °C, and the total CFU per unit area (Log CFU/cm^2^) of microtiter plate well were enumerated.

The determination of metabolic activity and total biomass were evaluated after washing each wells, according to Negri et al., 2016. The absorbance values 492 nm to XTT assay and 620 nm to CV assay were standardised per unit area of well (absorbance/cm^2^). The absorbance values of the negative controls wells were subtracted from the values of the test wells to account for any background absorbance [35].

For the analysis of matrix compounds, the matrix of different ages of maturation (24, 48 and 72 hours) was extracted using a protocol described by Capoci *et al.*, 2015 with some modifications. In brief, the biofilm samples were scraped from the 24-well plates, resuspended with PBS, and sonicated for 50 seconds at 30%, and then the suspension was vortexed vigorously. The suspension was centrifuged at 4000 ×g for 10 minutes, and the supernatant was filtered through a 0.22 μm nitrocellulose filter (Merck Millipore, Ireland) and stored at −20°C until analysis. Proteins, polysaccharides, eDNA and eRNA were measured by NanoDrop spectrophotometer (NanoDrop 2000 UV Vis Spectrophotometer, Thermo Scientific, Wilmington, DE, USA).

The morphological characteristic of *R. mucilaginosa* biofilm formation process (2, 24, 48 and 72 hours) was observed by SEM. For the SEM analysis, were performed according described previously. The samples were observed with a Quanta 250™ SEM (ThermoFisher) at 2000×magnification.

### 7.0 <u>In vivo</u> pathogenicity in model <u>Tenebrio molitor</u>

The evaluation of survival after infection of *Tenebrio molitor* larvae of approximately 100-200 mg using a total of 10 larvae per group. Three concentrations of inoculum with ATCC 64684 *R. mucilaginosa* were evaluated for standardization of the highest lethality inoculum: 1-3× 10³, 10^4^ and 10^5^ CFU in sterile PBS in aliquots of 5 μL were injected using a Hamilton syringe (701 N, 26’s gauge, 10 μL capacity), into the hemocoel, the second or third sternite visible above the legs and the ventral portion. Negative control included sterile PBS. The larvae were placed in sterile Petri dishes containing rearing diet and kept in darkness at 25 °C. Mortality was monitored once a day for 10 days. To establish larvae death, according to Souza et al., 2015 we visually verified melanization and response to physical stimuli by gently touching them.

With the standardized inoculum (1-3× 10^5^), we infected 10 larvae with each one of the clinical isolates in order to evaluate the virulence potential.

### 8.0 Ethics Statement

This study was conducted according to the Resolution 466/2012 of the National Health Council and was previously approved by the Ethics Committee for the Research Involving Humans of the State University of Maringá, Brazil under protocol numbers [COPEP-EMU n° 383979, CAEE resolution n° 17297713.2.0000.0104]. We obtained written and signed informed consent from the participant prior to study inclusion, who was an adult. The data collection, the oral mucosa examination, were performed according to Pieralisi et al., 2016.

### 9.0 Statistical Analysis

All tests were performed in triplicate, on three independent days. Data with a non-normal distribution were expressed as the mean ± standard deviation (SD) of at least three independent experiments. Significant differences among means were identified using the ANOVA test followed by Tukey’s multiple-comparison test. For *in vivo* pathogenicity, was using Kaplan–Meier survival plots, according to Souza et al., 2015. The data were analyzed using Prism 8.1 software (GraphPad, San Diego, CA, USA). Values of p < 0.05 were considered statistically significant.

## Results

Two clinical isolates obtained from saliva (CMRP3462) and sterile swab (CMRP3463) wiped in the center of the dorsal surface of the tongue collected were isolated and identified phenotypically by morphological plus biochemical aspects and confirmed as *R. mucilaginosa* by MALDI-TOF. In addition, the isolates were identified genotypically based on ITS region sequencing with following GenBank accession numbers CMRP3462 and CMRP3463 are MK453051.1 and MK453052.1. According to the phylogenetic analysis (Fig 1A), the isolates were located to the same clade of the type strain *R. mucilaginosa* CBS 316. The isolates were compared with clinical and environmental isolates from the other study [19] suggesting variability among the groups observed (Fig 1B).

**Figure 01:**
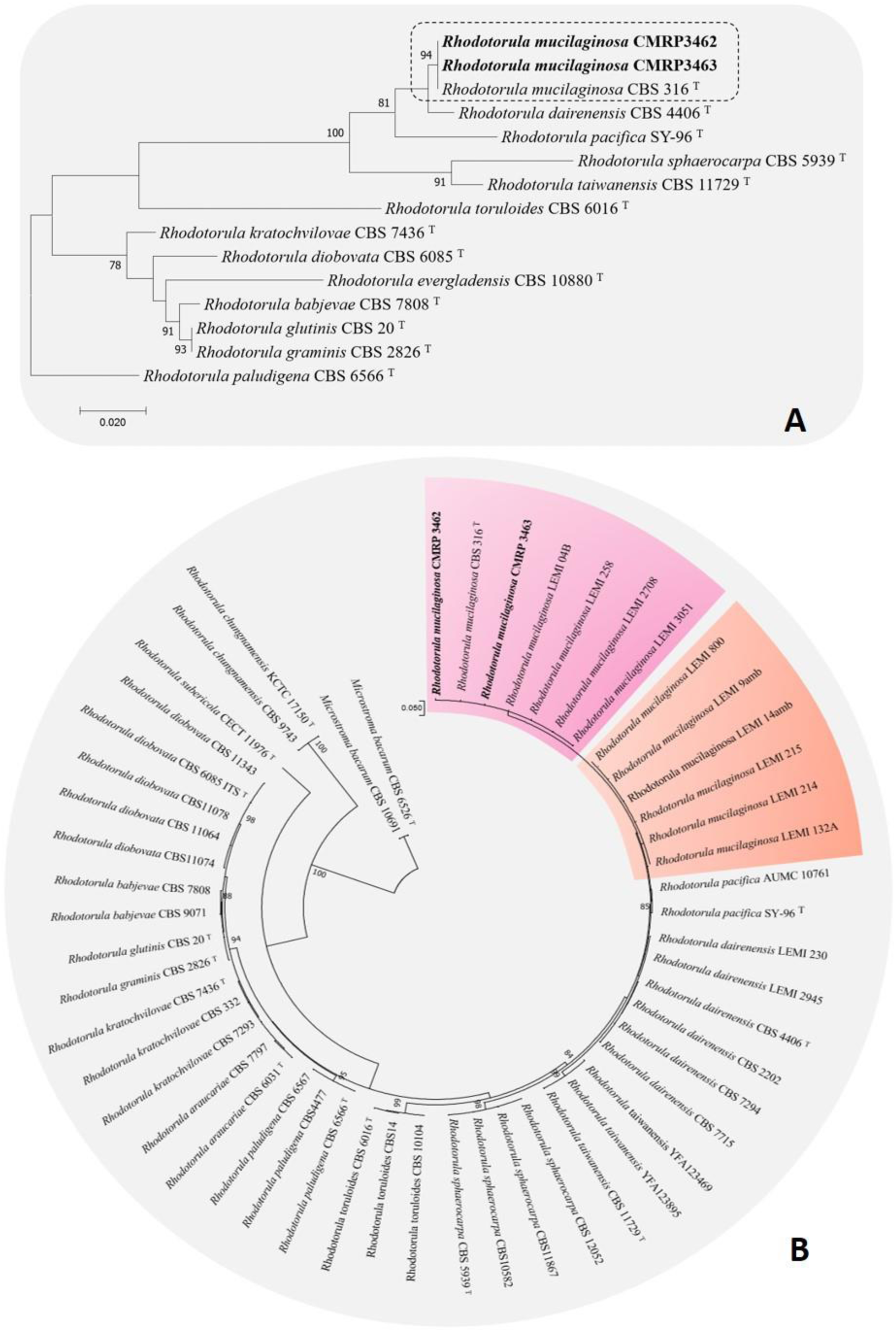
Phylogenetic analysis of *Rhodotorula mucilaginosa*, based on ITS sequences constructed with Maximum likelihood, based on the Tamura-Nei model + Gamma distribution (T92+G) implemented in MEGA v.7. Bootstrap support was calculated from 1000 replicates. (T) = type strain of the species. Bootstrap values > 80% were considered statistically significant. (A) Phylogenetic tree of *Rhodosporidium* clade, *Sporidiobolaceae* family*. Rhodotorula paludigena* CBS6566T was taken as outgroup. (B) Phylogenetic tree of *R. mucilaginosa* variability among clinical and environmental isolates of Nunes et al., 2013. (amb) = environmental lineages. *Microstroma bacarum* CBS 6526T and CBS10691 was taken as outgroup.

Observing cultures performed in SDA, we found orange-colored mucoid colonies (Fig 02 A), which grew within 48 hours at 25 °C. In the microscopic examination, the round blastoconidia, without the rudimentary formation of hyphae, were observed (Fig 02 B), as well as in the microculture (Fig 02 C) to confirm the micromorphological characteristics. Through China ink, it is possible to evidence *R. mucilaginosa* has a small polysaccharide capsule (Fig 02 D) and Calcofluor White revealed the cell wall of this yeast, which proves to be simple (Fig 02 E). Finally, with SEM, we observed with more clarity the round blastoconidia, in division (Fig 02 F).

**Fig 02.**
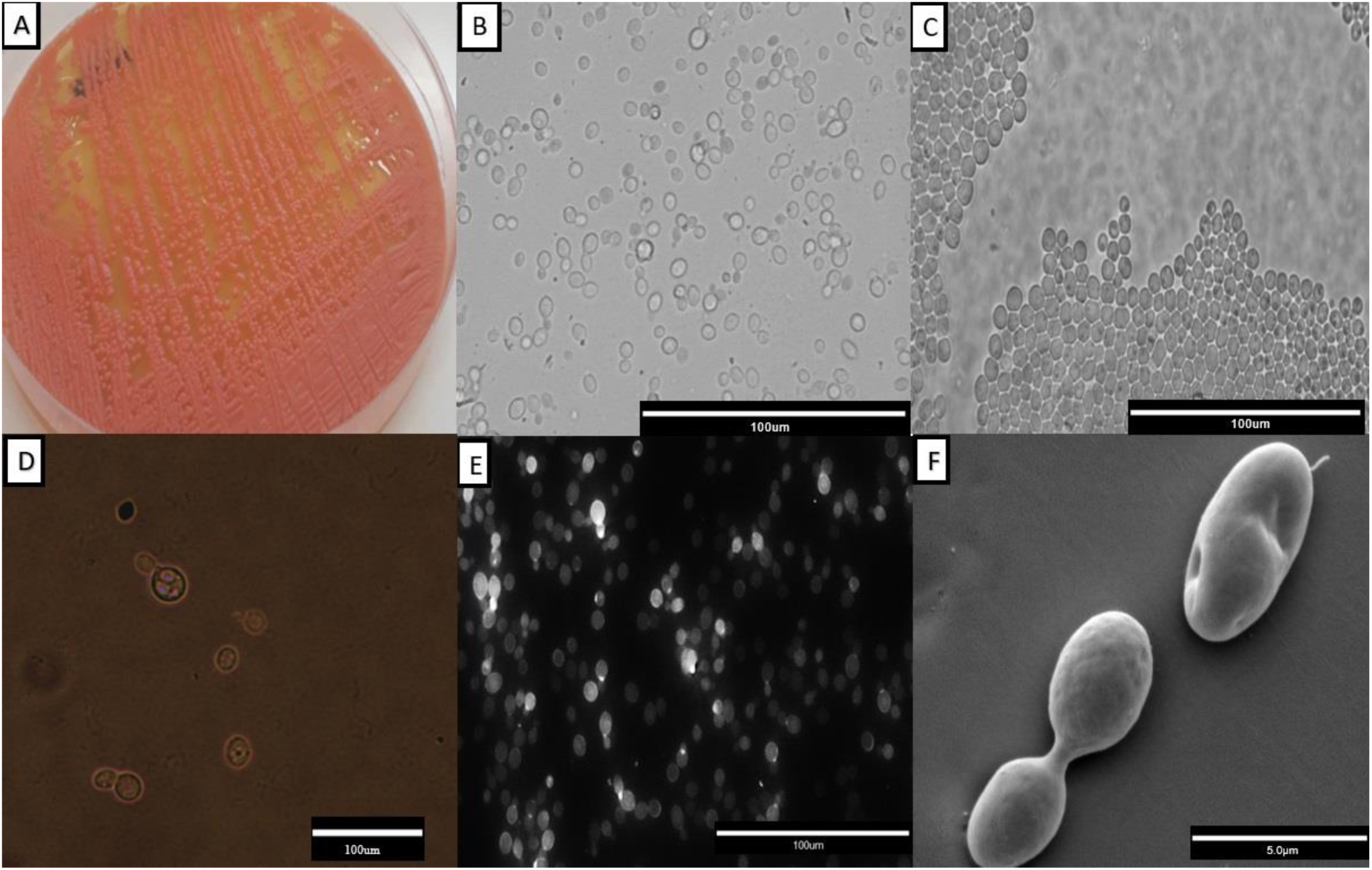
Representative morphological characterization of a *Rhodotorula mucilaginosa* isolate. In A, orange-colored mucoid colonies on Sabouraud Dextrose Agar; B, suspension of light field cells with a 40x magnification; C, characteristic microculture with rounded blastoconidias observed in a 40x magnification; D, polysaccharide capsule evidenced by China ink in 40x magnification; E, the cell wall evidenced by Calcofluor White; F, Scanning electron microscopy observed at 5000x magnification.

All isolates as clinical as ATCC 64684 were resistant to azoles (fluconazole, voriconazole, itraconazole), while for polienes, amphotericin B and nystatin, all isolates were sensitive (Table 01).

**Table 01.** Antifungal susceptibility profile of the clinical isolates and the reference strain was determined for the following antifungals: fluconazole, voriconazole, itraconazole, nystatin and amphotericin B, according to the guideline of the Clinical Laboratory Standards Institute (CLSI, 2008), and M27-A3 document.

| Strain | MIC*(µg/ml) |  |  |  |  |
| --- | --- | --- | --- | --- | --- |
|  | FLZ | VORI | ITRA | NYSTA | AMP-B |
| ATCC 64684 | 64(R) | 16(R) | 32(R) | 1(S) | 0,125(S) |
| CMRP3462 | 64(R) | 16(R) | 32(R) | 0,5(S) | 0,06(S) |
| CMRP3463 | 64(R) | 16(R) | 32(R) | 0,5(S) | 0,06(S) |

\*Criteria of interpretation (CLSI, 2008): FLZ, fluconazole ( $S \leq 8$ ; SDD 16–32; $R \geq 64$ ); VORI, voriconazole ( $S \leq 1$ ; SDD =2; $R \geq 16$ ); ITRA, itraconazole ( $S \leq 0,125$ ; SDD 0,25–0,5; $R \geq 32$ ); NYSTA, nystatin ( $S \leq 4$ ; SDD 8–32; $R \geq 64$ ); AMP-B, amphotericin B ( $R > 1$ ).

All the isolates of *R. mucilaginosa* showed the adhesion and biofilm formation abilities on the abiotic surface polistirene. In general, similar profiles among the isolates were observed (Figure 03). There was a significant increase of biofilm in number of cells until 48 hours of biofilm age, after this period there is a decrease of viability cells (Fig 03 A). On the other hand, metabolic activity and total biofilm biomass were different among the isolates, decreasing of 24 to 48 hours biofilm age (CMRP3463 and ATCC 64684) to metabolic activity (Fig 03 B) and increasing from 24 to 48 hours biofilm age (CMRP3462 and ATCC 64684) to total biofilm biomass (Fig 03 C). It is important to highlight that there was a decrease for all parameters analyzed after 48 hours of biofilm age.

**Fig 03.**
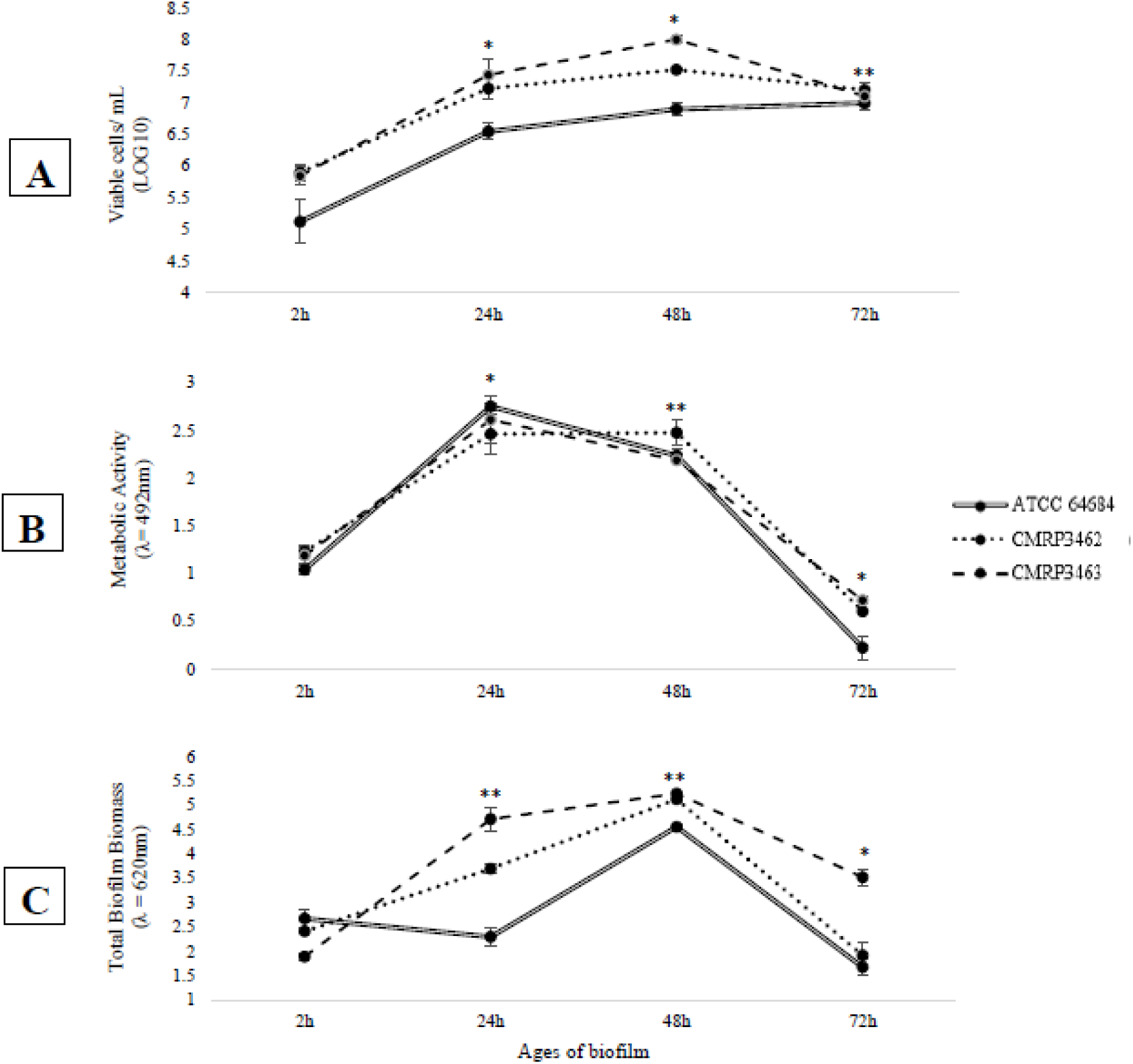
Adhesion capacity and biofilm formation, on polystyrene flat-bottom plates at different incubation times, of *Rhodotorula mucilaginosa*. A) Evaluation of cell viability by count of Colony Forming Units (CFU); B) Evaluation of metabolic activity by reduction of XTT; C) Evaluation of the production of extracellular matrix by Violet Crystal.*statistical difference in time among all isolates;**statistical difference over time for two isolates.

Analyzing each isolate, regard to adhesion (2 hours), ATCC 64684 strain showed the lowest cell viability by CFU with p<0.01 (Fig 03 A), the metabolic activity (XTT) was similar among the isolates (p>0.05) (Fig 03 B). However, in the evaluation of total biofilm biomass (CV), according to Fig 03 C, the CMRP3463 was the lowest total biofilm biomass (p<0.01). From 24 hours of biofilm formation, clinical isolates (CMRP3462 and CMRP3463) were significantly (p<0.01) higher than ATCC 64684 to viable cells in biofilm (Fig 03 A). All isolates presented a significant increase in relation 2 to 24 hours, with no statistical difference among isolates (Fig 03 B) to metabolic activity. The clinical isolates (CMRP3462 and CMRP3463) increase significantly (p<0.01) in the total biofilm biomass at 2 to 24 hours, mainly CMRP3463 (Fig 03 C). Finally, in the period of 48 to 72 hours, the clinical isolates (CMRP3462 and CMRP3463) showed a significant reduction of the number of cells viability (p<0.01). Further, there was a significant reduction for all isolates in metabolic activity and total biofilm biomass (p<0.01).

To the analysis of the extracellular matrix (ECM) of biofilms at different ages of maturation (24, 48 and 72 hours) eDNA, eRNA, proteins and polysaccharides were measured, as shown at Table 2. *R. mucilaginosa* were able to produce ECM in different ages of biofilm constituted of eDNA, eRNA, proteins and polysaccharides. These matrix compounds varied according to the time and strain. For ATCC 64684, there was a significant increase of eDNA between 48 and 72 hours, whereas for CMRP3462 there was a significant reduction between these same times. On the other hand, CMRP3463 showed no differences in the amount of eDNA in the biofilms of 24, 48 and 72 hours. When eRNA was evaluated, there was a significant increase between 24 and 48 hours, which remained constant at 72 hours for ATCC 64684. In relation to the clinical isolates CMRP3462 and CMRP3463, it was observed a contrary behavior, there is a greater amount of eRNA in 24 hours, while in 48 hours this amount is significantly lower and is maintained in 72 hours. For total proteins, there were no statistical differences among the isolates and the biofilm times evaluated. Finally, for polysaccharides we observed a significant increase at ATCC 64684 in 72 hours compared with the others ages of maturation of biofilm, while for the clinical isolates there were no statistical differences among the isolates and the biofilm times.

Through light microscopy and scanning electron techniques, can observe how occurs the biofilm formation by *R. mucilaginosa* which is shown by Fig 04, respectively. In all situations, *R. mucilaginosa* cells were without filamentation, in blastoconidia form, uniform size and oval shape. From the time of adhesion to 2 hours of incubation (Fig 04 A), can find a few scattered fungal cells or in small groups. After 24 and 48 hours, the yeasts were more clustered and in larger amounts, shown by Fig 04 B and C. Lastly, at 72 hours it is possible to observe that the fully established biofilm, with the confluence of *R. mucilaginosa* cells, shown by Fig 04 D. At scanning electron microscopy (Fig 04 D), still possible to see multiple layers of cells.

**Fig 04.**
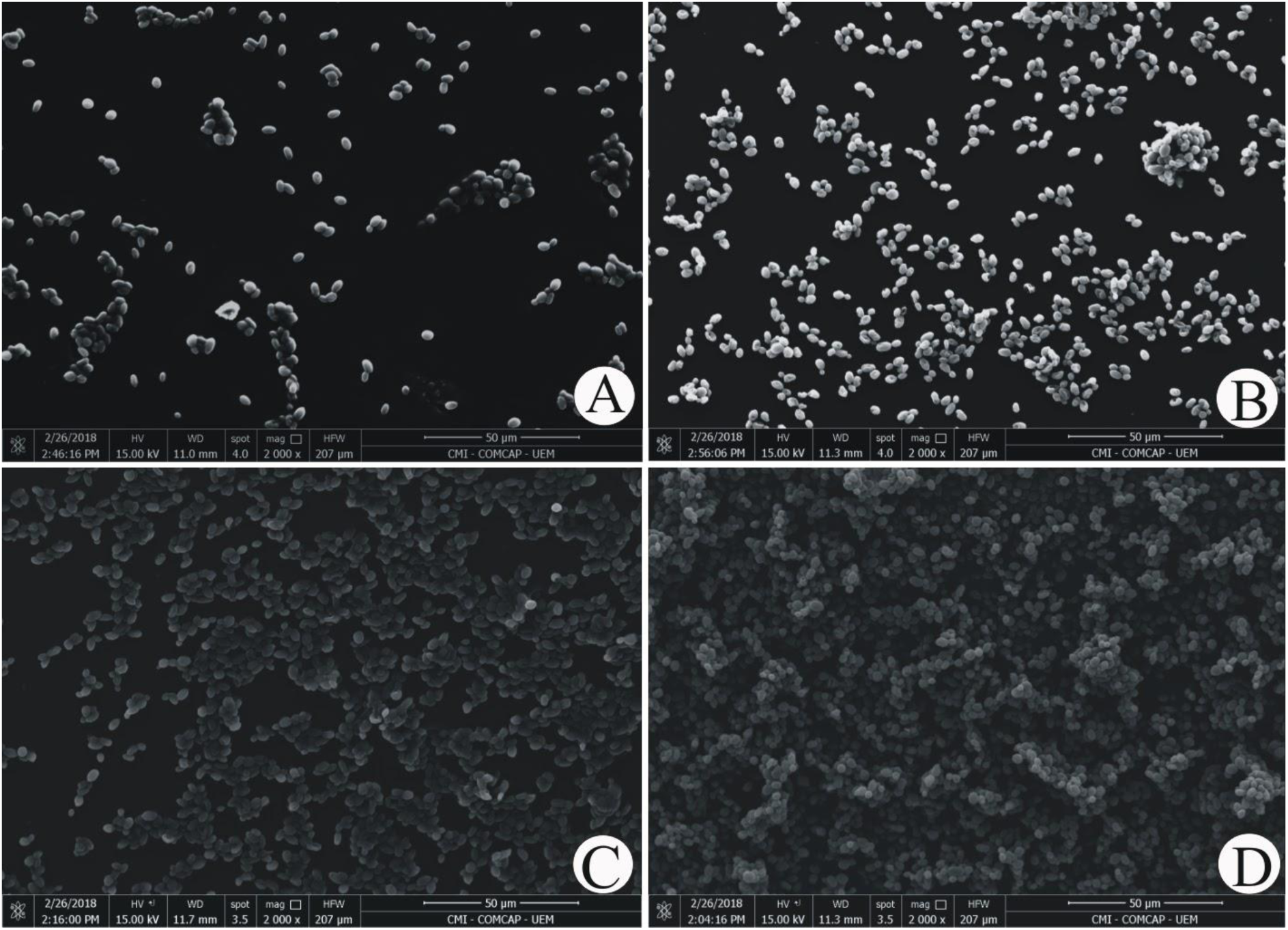
Illustrative images of ATCC 64684 *Rhodotorula mucilaginosa* adhesion and structure biofilms obtained by SEM, taken in a Quanta 250™ SEM (ThermoFisher,) magnification 2000×, shown differents ages of maturation (24, 48 and 72 hours). A) The adherence at 2 hours of incubation; B) The biofilm at 24 hours; C) The biofilm at 48 hours; D) The biofilm at 72 hours

In relation to the death curve, using *in vivo Tenebrio molitor* larvae model, three different inoculum concentrations for *R. mucilaginosa* ATCC 64684 (10^3^, 10^4^ and 10^5^) were evaluated. At the highest concentration (10^5^), we observed a great reduction in the survival percentage of the larvae in relation to the control and the other concentrations, in the first 24 hours. At the lowest concentrations (10^3^ and 10^4^), we found approximately 80% of survival at the end of the 10 days of evaluation, while in the highest concentration of the fungus, there was only a 40% survival (Fig 05). After defining the concentration 10^5^, we evaluated the clinical isolates, where we found that until the second day of evaluation, there was 15% death for CMRP3462 and 20% death for CMRP3463.

**Fig 05.**
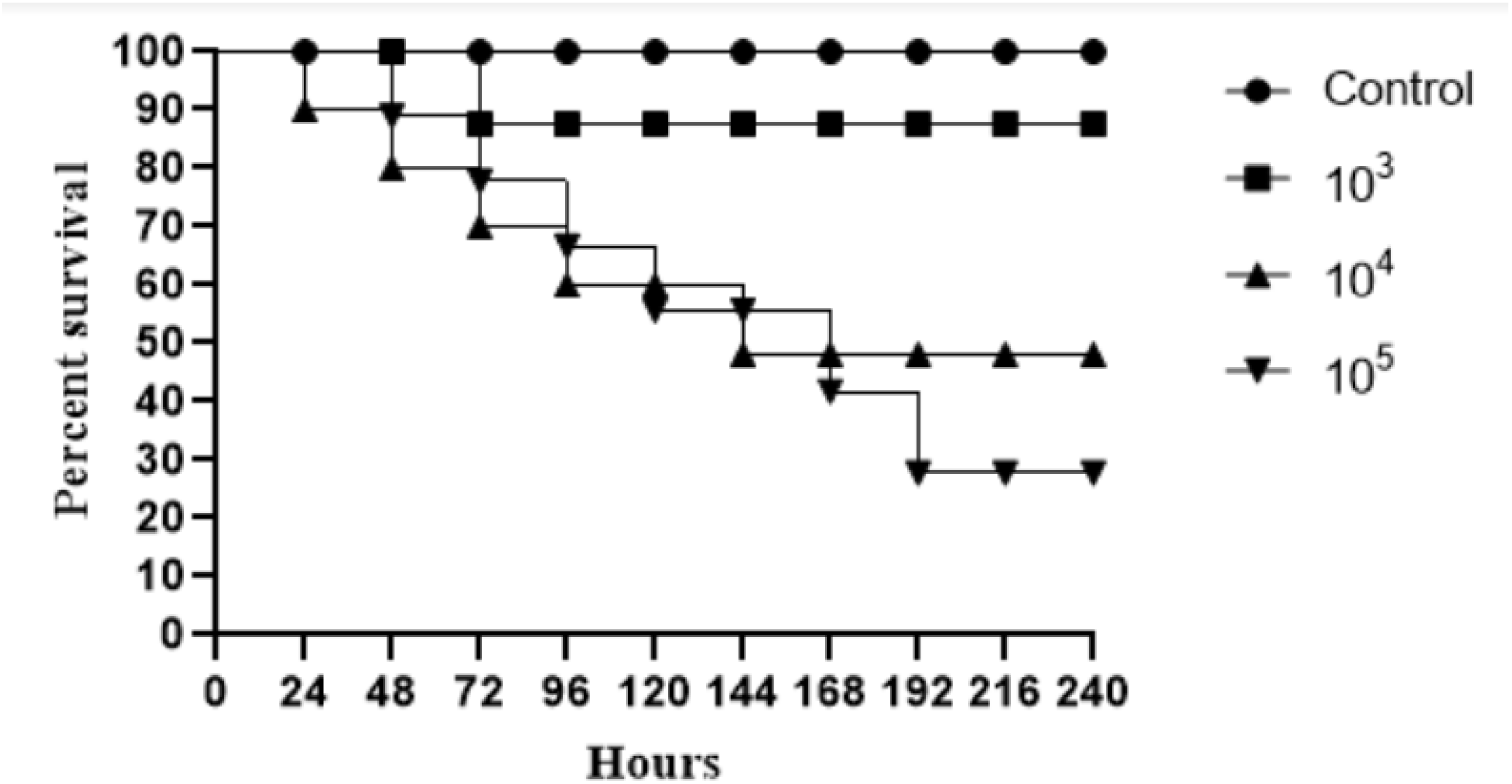
Survival curves of infected *Tenebrio molitor* with ATCC *Rhodotorula mucilaginosa*, for standardization. Groups of 10 larvae were infected with three fungal concentration. Negative control group the *T. molitor* larvae were injected just with PBS (without yeasts).

## Discussion

*Rhodotorula* spp. is a common saprophytic fungu**s**, being categorized as opportunistic and emerging pathogens recently [1]. An increase in the number of invasive infections by this yeast has been described in the last decades, with an overall mortality rate attributed aroud of 15% [10]. *R. mucilaginosa* is the most frequent species causing fungemia, which is responsible for up to 79% of infections, followed by *R. glutinis* (7.7%). These infections are often associated with the presence of CVCs or others implantable medical devices, and especially occur in immunocompromised individuals [6,8,12,23].

*Rhodotorula* is a polyphyletic nature group [33]. The majority species are environmental, however the family *Sporidiobolale* includes the type species *R. glutinis*, an opportunistic specie, and the emerging pathogen *R. mucilaginosa* (Fig 01). Besides *Rhodotorula* species are distributed among family of *Pucciniomycotina phylum* [33].

Current study included two clinical isolates of *R. mucilaginosa* obtained from a CKD patient living in south of Brazil, phylogenetic analysis was performed with sequences deposited by Nunes et al., 2013, who evaluated isolates from 51 clinical and eight environmental isolates recovered from 14 different Brazilian hospitals from 1995 to 2010. Our goal was to correlate our samples with the bank of those authors (complementary material) and no differences were found between clinical isolates from different human sites, or between clinical and environmental isolates, neither geographic differences, these data reinforce the ubiquitous character of *Rhodotorula* spp. Nevertheless, as only ITS regions were evaluated, more in-depth studies, such as multi-locus region analyzes would be interesting in order to confirm this possibility [38].

Regarding to micromorphological characterization we draw attention to a unique pattern of images displayed by *R. mucilaginosa*, found in all methodologies for both as in their planktonic forms (Fig 02) as in formed biofilms (Fig 04). Figure 02 shows the coral-red-color colonies characteristic of the genus *Rhodotorula*, rounded blastoconidia, with absence of filamentation and presence of polysaccharide capsule, these characteristics are similar to those described by Gan *et al.*, 2017 and Kitazawa *et al.*, 2018.

The low morphological variation could also hinder the laboratory diagnosis in routine laboratories. Important to note that, in current study clinical isolates of *R. mucilaginosa* were correctly identified by morphological characterization together with the biochemical identification tests (assimilation and fermentation). These are simple and inexpensive tests, known by all laboratories and they were sufficient to identify this species, confirmed through MALDI-TOF and later through molecular techniques. In fact, macromorphology, especially the typical color exhibited by their colonies on SDA and micromorphology on cornmeal-Tween 80 agar are considered key characteristics for the presumptive diagnosis of this genus [19].

At the same time, is important highlight alone, these observation can misendificate *Rhodotorula* with *Cryptococcus*, as beside the morphological similarity, both microorganisms are urease positive, are not capable of fermentation and can assimile glucose, maltose, sucrose, galactose, xylose, raffinose and trehalose [26]. Despite recently, Yockey et al., 2019 suggested that *R. mucilaginosa* cells possess differences in signaling pathways, cell wall composition and that their membranes are more susceptible to perturbations than those of *C. neoformans*, similarities are observed in fungi morphology as well as in relation to the clinical aspects of the diseases caused by these two genus. Therefore, in some cases modern techniques are required for differentiate these microorganisms, such as the reporting of George *et al*., 2016. These authors found a CKD patient with an unhealed lesion on the right elbow, multiple biopsies suggested cryptococcal infection with necrotizing granulomas. Although, a panfungal PCR on a skin biopsy identified as *Rhodotorula* sp. Summarizing, the laboratory supplies an important and definitive approach for differential diagnosis between these important diseases. The correct diagnosis becomes indispensable, as according to Ioannou *et al.*, 2018, 16.2% of the systemic infections by *Rhodotorula* sp. attacked the CNS, where this species is the second most common fungal agent, affecting especially immunocompromised individuals [1].

In this kind of patient is historically predisposed to CNS infections by *Cryptococcus* sp. which despite the similarity with *Rhodotorula* sp. above discussed, the antifungal of choice are different, as fluconazole is indicate for *Cryptococcus* sp. while yeasts of the genus *Rhodotorula* are resistant to this antifungal class [43]. It is important to note that yeasts belonging to *Rhodotorula* genus have a high level of fluconazole resistance, a greater tolerance for itraconazole and susceptibility for amphotericin B [44, 45]. In agreement, we found in the antifungal susceptibility profile of *R. mucilaginosa* isolates resistant to azoles and sensitive to the other antifungals evaluated (Table 01). Recently, Wang *et al.*, 2018 have demonstrated that patients undergoing treatment with echinocandins addressed to other fungal infections have been reported to have fungal infections caused by *Rhodotorula* spp. since this yeasts are less susceptible to echinocandins due to the absence of 1,3-β-D-glucan in their cell walls.

In order to explain why this genus have increasing in disseminated and severe human infections, besides the resistance to antifungals, it is possible that this yeast have putative virulence factors. However, there are few studies in the literature that evaluate the virulence of *R. mucilaginosa* [19, 47] thus, much information are still unknown.

Considering the lack of filamentation capacity, it makes important the search to understand what would be the mechanism of pathogenicity. Biofilm production ability is one of the first suspicion due the the association of this genus with the use of CVC [1]. However, in contrast to the extensive literature dealing with biofilms of *Candida* sp. [18,20,35], few studies on the biofilm produced by isolates of medical interest of *R. mucilaginosa* are available. This lack of knowledge deserves concern, since serious and fatal infections due this specie have been related to the formation of biofilms on medical devices [11, 12].

The first study to evaluate the biofilm formation capacity of different *Rhodotorula* species [19] found differences between clinical and environmental isolates, through crystal violet staining only in 48 hours biofilms. Among the main results, *R. mucilaginosa* were classified as average biofilm producers, according to the classification scale for biofilm formation, adopted by the authors, which is in agreement to our results on biomass quantification.

The present study is the first one aiming to characterize the biofilm production by *R. mucilaginosa* on different ages (24, 48 and 72 hours), then we employed the classic methods used in studies with *Candida* sp. biofilms [35, 48], that are CFU, CV, XTT and microscope, addressing to *R. mucilaginosa*. These results are presented on Fig 03 and Table 2, and complemented by the quantification of the extracellular matrix components determined on the same times (Table 02). It was possible to observe a formation and organization of the biofilm over time, with apex in 48 hours and probable dispersion at 72 hours, when there is decay of the ECM, metabolic activity and CFU. Although there is a general trend among isolates, it is possible observe a weak difference in behavior at some specific times, suggesting characteristics isolate-dependent. In view of this scenario, we have inferred that ECM from *R. mucilaginosa* is organized differently from other yeasts, like *C. albicans* as are not view filaments or other specialized structures, however its architecture is similar to other pathogens unable of filamentation such as *C. glabrata* and *Cryptococcus* spp. [49].

**Table 02.**
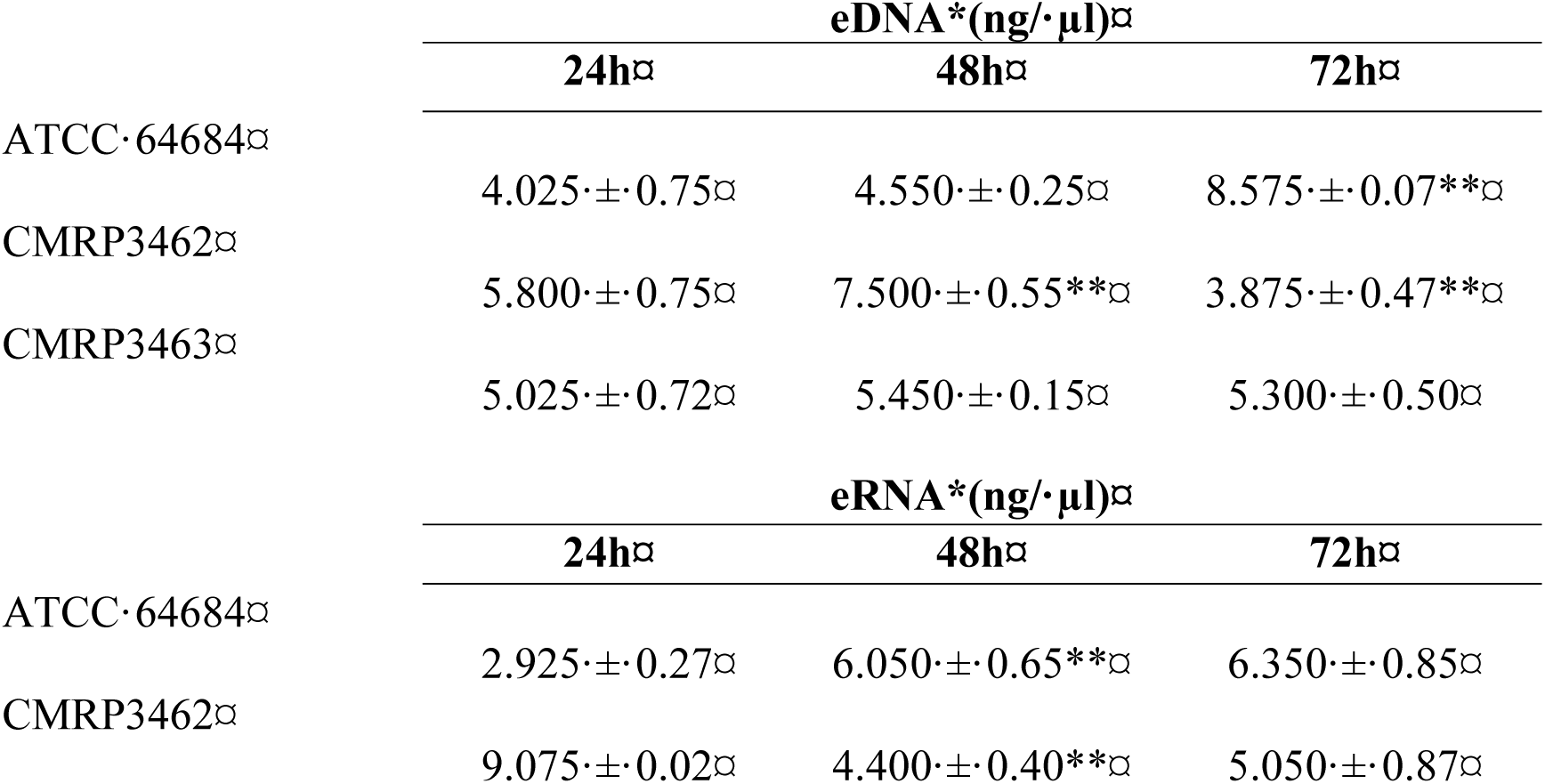
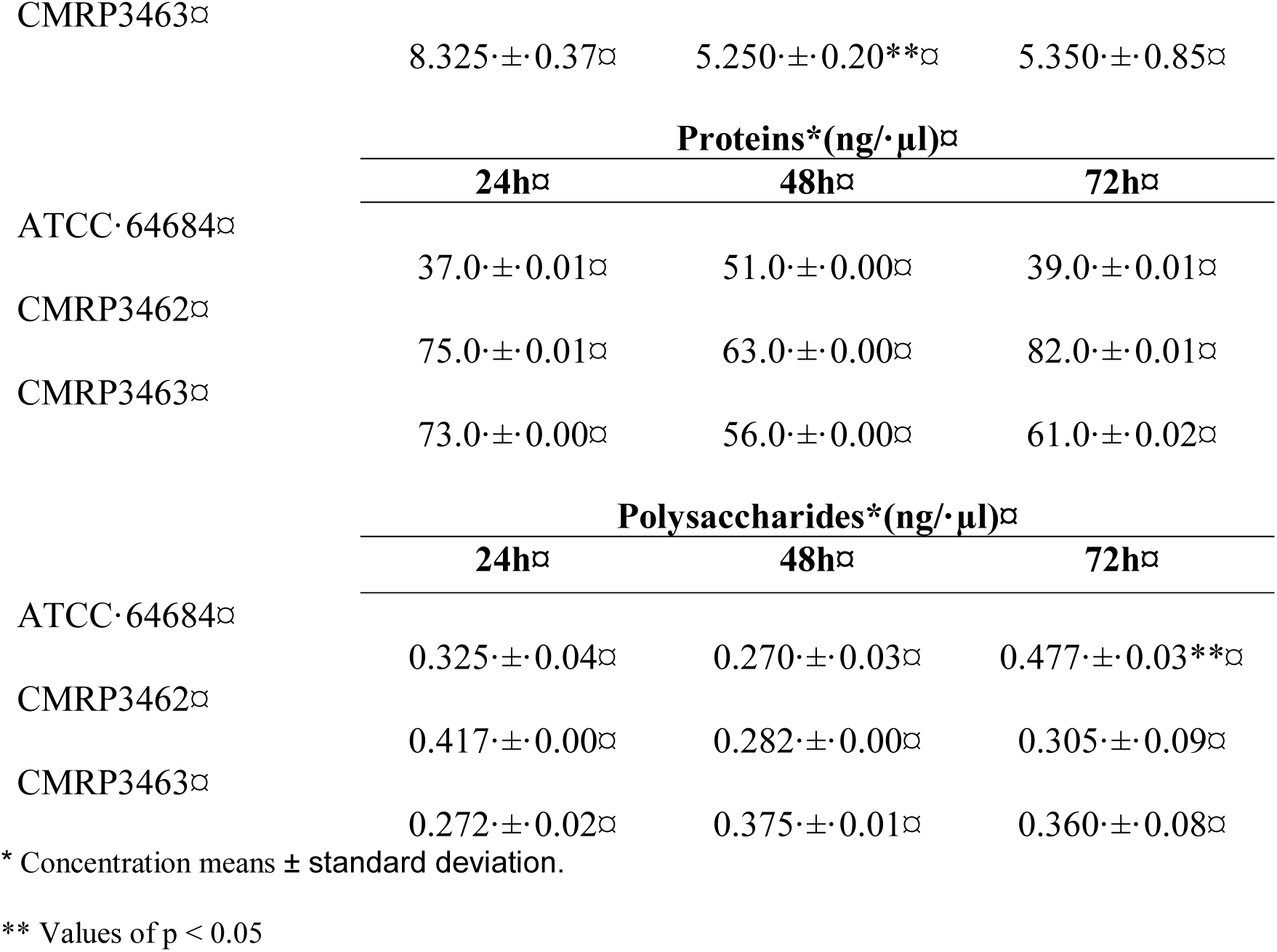
Quantification of extracellular DNA, extracellular RNA, proteins and polysaccharides for biofilm matrix analysis performed by NanoDrop spectrophotometer (NanoDrop 2000 UV Vis Spectrophotometer, Thermo Scientific, Wilmington, DE, USA).

On the other hand, seeing the biofilm architecture shown in Fig 04, we have hypothesized would be occurring with *R. mucilaginosa* a similar event to what occurs to the biofilm of *Staphylococcus aureus*, where there is the detachment of microcolonies of biofilm and the rolling of these biofilm microcolonies [50]. Thus, would be possible, attached cells, as well as part of the extracellular matrix have been removed during the analysis steps. According Rupp *et al.*, 2005 this is important mechanism, mainly for non motile microorganisms, as the controlled dispersal along surfaces in the protected biofilm state [50]. Therefore, even so few information from Fig 03 and Table 2, biofilm by *R. mucilaginosa* would be important as infection source since detached yeast from biofilm would be in the same way being carried. Thus, is reasonable to assume that detached yeasts from mature biofilm on CVC could reach host cells and other sites of the human body by dissemination, as it has been previously described for *C. albicans* [51].

In fact, recent studies, with other pathogenic fungi, have shown the cells that detach themselves from a biofilm have a greater association with mortality, compared to microorganisms in their planktonic form. More than 65% of human infections involve the formation of biofilms, which is related to the growing number of immunocompromised patients. In addition, more than 500,000 deaths per year are caused by biofilm-associated infections [21, 52]. Our results corroborate this idea, since according to Fig 03, we found a high value of viable cells in biofilms, for all isolates of *R. mucilaginosa*, similar to that occurring in *C. albicans*.

In addition, we saw a peak of metabolic activity in 24 hours, and soon afterwards a decrease of this activity, being that in 72 hours, these cells were probably “dormant” [35]. Metabolically “dormant” yeast cells are also known as persistent cells, which originate stochastically as phenotypic variants within biofilms [20]. According to Kojic et al., 2004, persistent biofilm cells represent an important mechanism of resistance, and the eradication of a biofilm usually requires the administration of toxic concentrations of antimicrobials, and the recommended treatment includes removal of the contaminated device. These ours findings could justify the association between *R. mucilaginosa* infections and biofilm formation.

SEM images reinforce our results on the biofilms evaluation, we observe that in the course of time, there was an increase in the number of cells and their organization. Unlike *C. albicans, C. parapsilosis or C. tropicalis*., we did not find filaments that give the structure for a complex biofilm, however we found several layers of cells, as well as described by Nunes et al., 2013 [19, 54]. With these variables, we infer that the maturity of *R. mucilaginosa* biofilm occurs in 48 hours due to stability and uniformity, confirmed mainly by microscopy images (Fig 04).

In view of the weak arguments that would justify the increase of *R. mucilaginosa* infections, we performed an *in vivo* infection on the invertebrate host *Tenebrio molitor*, an important tool to evaluate virulence of clinical pathogenic yeast strains [37]. We have observed the development of this larvae and its resistance to external stimulus or situations, then in our opinion it is a model of competent host, and it was fundamental since other *in vivo* studies found in the literature, were made with immunosuppressed animal [47]. Here, *T. molitor* larvae were used, for the first time, with *R. mucilaginosa*, in order to evaluate the pathogenicity of this species. Surprisingly, the survival curves obtained (Fig 05) were similar to those found for *Cryptococcus neoformans* [37], a recognized human pathogen, which can be misidentified with *Rhodotorula* spp. regarding clinical and laboratory aspects [42]. Increasing concentrations of inoculum (10^5^) of this yeast resulted in high mortality rates, confirming the efficiency of the method to evaluate the virulence of pathogenic yeasts and showing, for the first time, the pathogenic potential of *R. mucilaginosa.* Despite being the highest fungal concentration tested in this study, these results suggest high risks of infection and lethality also in humans. Summarizing, CKD patients are usually colonized by *Rhodotorula* spp. [5, 14]; moreover their immunosuppressed state [17] and considering the potential of virulence of this microorganism demonstrated in current study, we can infer CKD patients are critical risk group for disseminated infection by *R. mucilaginosa*.

## Conclusion

In this way, we infer that colonization of chronic renal patients by *R. mucilaginosa* offers a high risk of serious infection since this yeast showed highly pathogenicity for *in vivo* model, suggesting high risks of infection and lethality. Besides that, it is able to form biofilm on the surfaces of the medical devices, and apparently, the attached cells, as well as part of the extracellular matrix are removed and would fall into the circulatory chain being an important source of systemic infection. In addition, it is highly resistant to conventional antifungal agents, even antifungal of the last generation. We also emphasize that the correct identification of yeast is the main means for an efficient treatment.

## Acknowledgments

This study was supported by Coordenação de Aperfeiçoamento de Pessoal de Nível Superior (CAPES), Conselho Nacional de Desenvolvimento Científico e Tecnológico (CNPq), Fundação de Amparo à Pesquisa do Estado do Paraná (Fundação Araucária) and Financiadora de Estudos e Projetos (FINEP/COMCAP).

The authors would like to thanks to Laboratory of electron microscopy and microanalysis, Universidade Estadual de Londrina (UEL/SETI).

